# Bronchopulmonary Dysplasia with Pulmonary Hypertension Associates with Loss of Semaphorin Signaling and Functional Decrease in FOXF1 Expression

**DOI:** 10.1101/2024.08.27.609955

**Authors:** Shawyon P. Shirazi, Nicholas M. Negretti, Christopher S. Jetter, Alexandria L. Sharkey, Shriya Garg, Meghan E. Kapp, Devan Wilkins, Gabrielle Fortier, Saahithi Mallapragada, Nicholas E. Banovich, Christopher V. E. Wright, David B. Frank, Jonathan A. Kropski, Jennifer M. S. Sucre

## Abstract

Lung injury in preterm infants leads to structural and functional respiratory deficits, with a risk for bronchopulmonary dysplasia (BPD) that in its most severe form is accompanied by pulmonary hypertension (PH). To examine cellular and molecular dynamics driving evolving BPD in humans, we performed single-cell RNA sequencing of preterm infant lungs in early stages of BPD and BPD+PH compared to term infants. Analysis of the endothelium revealed a unique aberrant capillary cell-state primarily in BPD+PH marked by *ANKRD1* expression. Predictive signaling analysis identified deficits in the semaphorin guidance-cue signaling pathway and decreased expression of pro-angiogenic transcription factor *FOXF1* within the alveolar parenchyma in neonatal lung samples with BPD/BPD+PH. Loss of semaphorin signaling was replicated in a murine BPD model and in humans with alveolar capillary dysplasia (ACDMPV), suggesting a mechanistic link between the developmental programs underlying BPD and ACDMPV and a critical role for semaphorin signaling in normal lung development.

## Introduction

In infants born preterm, abnormal lung development leading to chronic lung disease is a leading complication, resulting in bronchopulmonary dysplasia (BPD)^1^. In its most severe form, BPD is associated with life-threatening pulmonary hypertension (BPD+PH), which is thought to be caused by abnormal development of the pulmonary vasculature, with decreased numbers of alveolar-capillary units formed for gas exchange^2,3^. Despite increasing awareness of the environmental and perinatal risk factors that increase the risk of BPD, there are very few disease-modifying therapies, and our understanding of the molecular mechanisms and potential therapeutic targets remains limited.

The advent of single-cell multiomics technology as applied to human lung samples has enabled an increasingly nuanced understanding of the cell populations and cell-states under homeostatic and diseased conditions in adults^4–6^. However, because neonatal biopsies are infrequently performed and infant autopsies are rare in general, lung tissue from preterm infants and healthy neonates is relatively under-represented in most of the currently available large sequencing series. Here we report single-cell transcriptomic findings from former preterm infants with evolving BPD, and infants with both BPD and PH, compared to term infants at approximately the same corrected gestational age as the preterm infants.

While we anticipate that this transcriptional dataset will be a resource for future work in neonatal lung injury and BPD in general, we were particularly interested in the endothelial cell populations in infants with BPD+PH and how they might differ from corrected gestational age-matched controls and patients with BPD at a similar age. In these sequencing data, we identified a capillary endothelial cell-state highly specific to patients with BPD+PH and spatially localized these capillary cells to the distal alveolar parenchyma, and not in more proximal tissue areas. Further, we employed predictive signaling analysis to generate hypotheses about the role of semaphorin guidance-cue signaling during alveolar formation, a pathway we identified as deficient in both BPD and BPD+PH patients transcriptomically, and then spatially localized to the developing alveolar regions. Our parallel identification of semaphorin signaling deficits in alveolar epithelial and endothelial cells in an established mouse model of BPD suggests that this pathway is central to normal alveologenesis. Further, we find evidence of dysregulated semaphorin signaling in patients with alveolar capillary dysplasia with misaligned pulmonary veins (ACDMPV), implicating semaphorin signaling as a potential mechanism driving abnormal lung development after preterm birth.

## Results

To examine the cellular dynamics and candidate mediators of neonatal lung injury and the development of severe BPD, we used the 10x Genomics Chromium Platform to perform single-cell RNA sequencing (scRNAseq) of human neonatal lungs at varying stages of lung injury and chronic lung disease: one acute preterm infant with lung injury, two BPD samples at term corrected gestational age (CGA), two BPD+PH at term CGA, and two term infants in the first three weeks of life for age-matched comparison (**Fig. 1A**, **Supplementary Table 1**). After filtering and quality control to remove doublets and ambient RNA (see materials and methods), we recovered 43,607 single-cell transcriptomes with a range of 2,199 to 8,339 cells per subject, a median of 4,128 unique molecular identifiers (UMI) (interquartile range 1,188 – 4,670) per cell, and a median of 1,795 genes (interquartile range 728 – 1,944) per cell (**Fig. 1A-B**). Unbiased clustering with cell-type annotation informed by prior human, fetal, and neonatal studies identified seven epithelial, seven endothelial, seven mesenchymal, and twelve immune-cell clusters (**Fig. 1B-G**)^4–12^. To specifically focus on potential vascular cell differences in BPD patients with and without PH, the endothelial cluster was extracted, re-embedded, and separately analyzed to identify cell-type-specific differences across homeostasis and disease. These sequencing data are available through an interactive data explorer at www.sucrelab.org/lungcells.

**Fig. 1:**
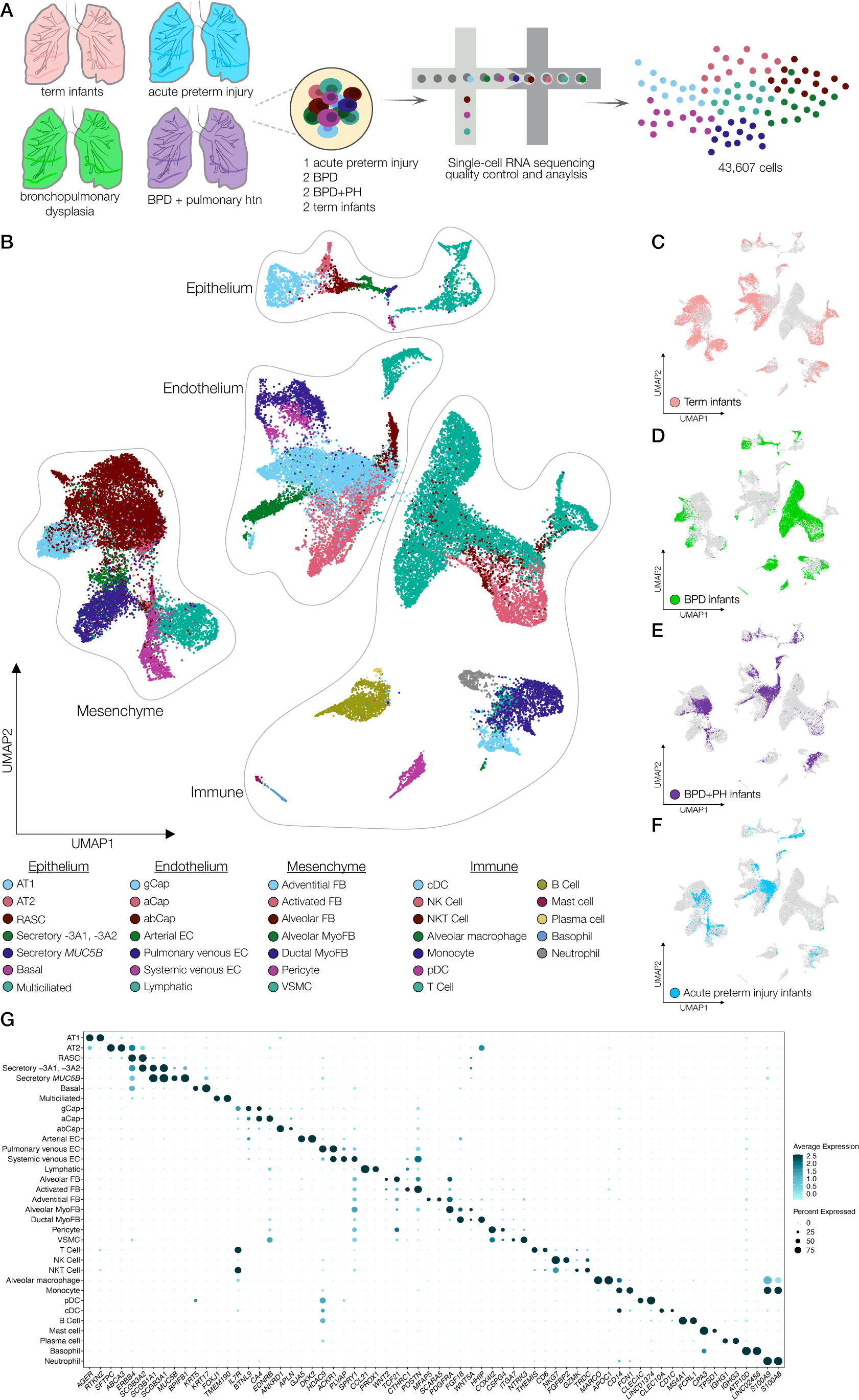
Analysis of neonatal human lung single-cell transcriptomics identifies distinct populations of epithelial, endothelial, mesenchymal, and immune cells. A) Experimental design overview for single-cell sequencing of two term control infants, one acute neonatal lung injury, two infants with severe bronchopulmonary dysplasia, and two patients with severe bronchopulmonary dysplasia and pulmonary hypertension. B) Uniform Manifold Approximation and Projection (UMAP) embedding of the neonatal human lung dataset (n = 43,607 cells). Cells labeled by lineage and by cell type. C-F) Breakdown of the dataset’s UMAP embedding by disease-state condition annotation. G) Dot plot of the hallmark-identifying genes for each cell cluster. Dot size corresponds to the percentage of cells within a cluster expressing a certain gene. Color intensity corresponds to the relative average expression level.

### Bronchopulmonary dysplasia with pulmonary hypertension is associated with a specific population of capillary endothelial cells

Analysis and re-embedding of the endothelial cells identified a population primarily obtained from BPD+PH patients with transcriptional hallmarks common to both alveolar capillaries (aCaps, CAP2) and general capillaries (gCaps, CAP1) (**Fig. 2A-B**, **S1A**). This group of 647 cells is marked by high expression of *ANKRD1*, a gene which is nearly absent among all other endothelial cell clusters, with limited detection in alveolar type 1 (AT1) cells (**Fig. 2C**). This aberrant capillary cell-state (abCap) expresses some aCap-associated genes, e.g. *APLN*, *EMCN*, *S100A4*, but has markedly reduced expression of aCap hallmark genes *EDNRB* and *SOSTDC1* when compared to other aCaps. Similarly, abCaps express some genes associated with gCaps, e.g., *PTPRB* and *EDN1*, but lack expression of canonical gCap genes *FCN3*, *IL7R*, and *BTNL9* (**Fig. 2D**). The abCaps also express certain pan-capillary markers *PDGFB*, *RGCC*, and *ADGRL2*, but not *CA4*, *CD36*, and *AFF3*. In addition to the YAP/TAZ signaling target *ANKRD1*, this population expresses relatively high levels of other YAP targets *CCN1* (*CYR61*), *CCN2* (*CTGF*), *SERPINE1*, and *CRIM1*, compared to other endothelial cells (**Fig. 2D**). Notably, the abCaps present a gene-expression profile similar to previously described endothelial tip cells^13,14^, a population present in early lung development that directionally guides the growth and migration of capillary endothelium. Relative to other endothelial cells, abCaps have increased expression of tip cell hallmarks *ESM1*, *ANGPT2*, *APLN*, *MCAM*, *COTL1*, but lower expression of tip cell–associated receptors such as *KDR*, *FLT4*, *UNC5B*, and *NRP1,* relative to other endothelial cells (**Fig. 2E**).

**Fig. 2:**
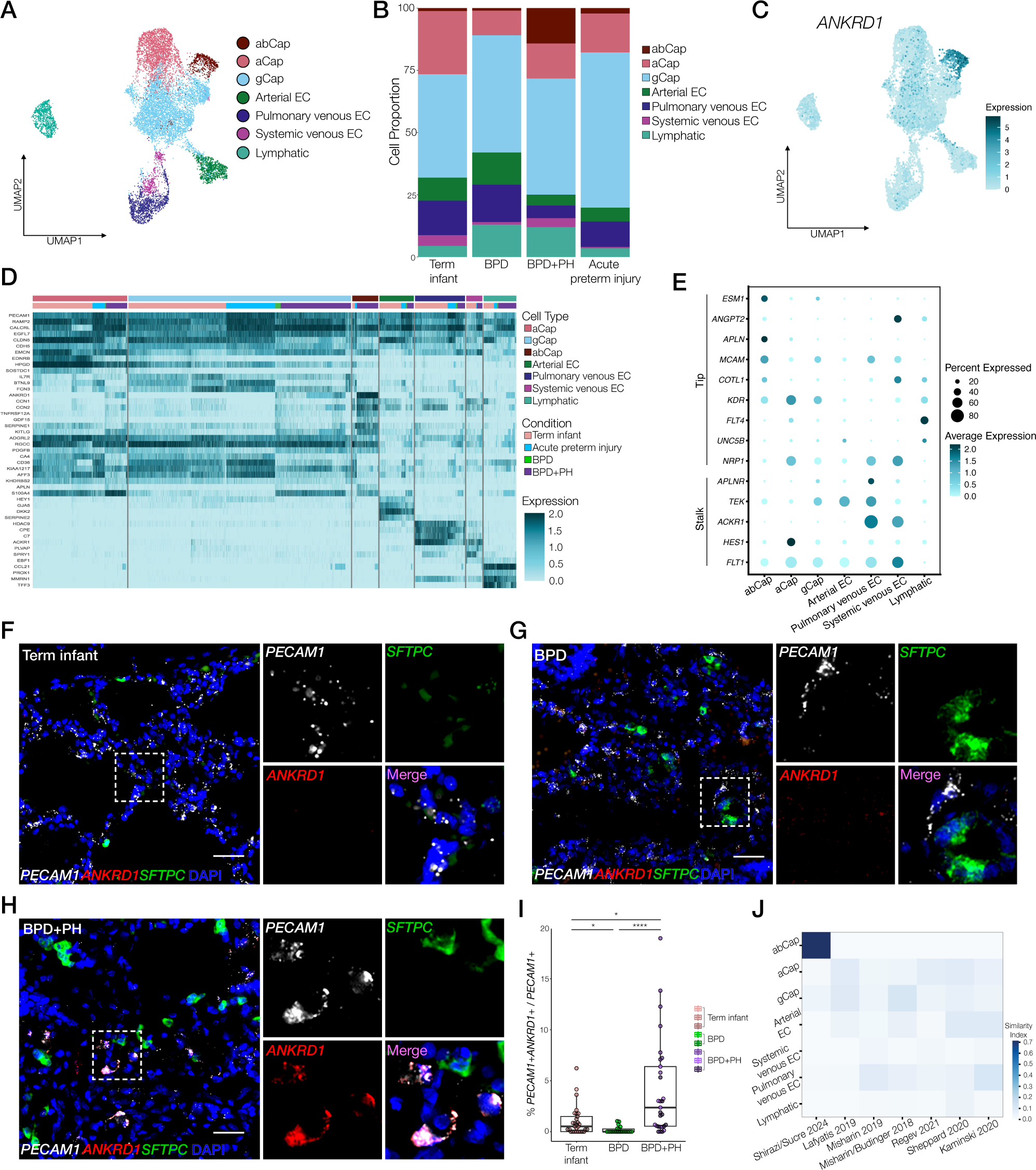
Analysis of the endothelium reveals a cluster of aberrant capillary cells (abCaps) enriched in patients with BPD+PH. A) UMAP embedding of the neonatal human lung endothelial cells (n = 11,816) labeled by cell type. B) Stacked area plot of cellular proportions across the four disease-state conditions. C) UMAP of the expression level of *ANKRD1*, the highest expressing gene in the abCaps. D) Heatmap of hallmark genes of the seven identified endothelial cell type clusters, demonstrating a distinct expression profile for abCaps when compared with aCaps and gCaps. The cell types are subdivided by disease condition. Maximum expression values are clipped at 2. E) Dot plot of the hallmark genes for endothelial tip and stalk cells. Dot size corresponds to the percentage of cells within a cell type cluster expressing a certain gene. Color intensity increases with relative average expression level. F-H) RNA ISH for *ANKRD1* (red), *PECAM1* (white), and *SFTPC* (green) demonstrating co-localization of *ANKRD1* within *PECAM1*+ cells in BPD+PH patients in the alveolar parenchyma. Scale bar = 30 μm. I) RNA ISH quantification using HALO of the percentage of *PECAM1*+ cells that are also *ANKRD1*+ over the total amount of *PECAM1*+ cells (*PECAM1*+*ANKRD1*+/ all *PECAM1*+) across disease condition, * *p-*value<0.05, **** *p*-value<0.0001, Kruskal-Wallace one-way ANOVA. J) Heatmap showing relative homology from top 6 datasets identified by SCimiliarity, with intensity of color indicating greater similarity, abCap cell included in upper right at positive control.

To spatially validate and localize the abCap cell-state, we performed RNA *in situ* hybridization (RNA ISH) using formalin-fixed paraffin-embedded (FFPE) lung tissue blocks from preterm infants with BPD with and without PH, as well as from term infant control lungs (**Fig. 2F-H**). These FFPE tissues are from an expanded set of samples, independent of those interrogated by scRNAseq (**Supplementary Table 2**). In lung tissue from patients with BPD+PH, *ANKRD1*+/*PECAM1*+ endothelial cells were localized to the alveolar parenchyma, near *SFTPC*+ alveolar type 2 (AT2) cells, consistent with their expression of capillary hallmark genes. Quantification demonstrated a marked enrichment of this population in patients with BPD+PH when compared with age-matched BPD patients and normal term infant controls of approximately the same corrected gestational age **(Fig. 2I**).

### The abCap endothelial cell state does not transcriptionally resemble endothelial cells previously described in different human lung diseases

To understand abCaps in the context of previously reported atlases of human pulmonary endothelium under normal and disease conditions, we performed analysis with SCimilarity^15^, a computational tool that allows comparison of a given cell population with cell populations from nearly 400 previously published human datasets, including 41 from the lung. Comparing abCaps across the 1.12-million endothelial cells in the 22.7-million cell corpus of SCimilarity, we selected the top 100 human samples (contained in 34 source studies) with the highest SCimilarity score to our BPD+PH capillary cells. Of these 34 studies, we excluded seven studies of iPSCs, 14 that did not contain lung tissue, two with samples from patients in the 7^th^ – 8^th^ decade of life, two with a sample size < 2, and two with inaccessible scRNAseq data files. This subtraction left six studies with 178 samples for comparison (**Fig. S1B**). We compared the abCap BPD+PH endothelial cell state to the endothelial cells of healthy controls and five disease states – COVID, LAM, IPF, SSc-ILD, and PAH. Using the genes most specific for abCaps, we created a gene-module score to identify cells with expression patterns characteristic of abCaps (see methods for details). Among these six studies, fewer than 12% of cells in each endothelial sub-population were showed abCap similarity (**Fig. 2J**). As a positive control that the selected module score is inherently selective of the abCap population, applying this metric to our entire dataset confirmed that the cells containing the genes with the highest gene-module score were within the BPD+PH associated abCap population.

### Analyzing putative ligand-receptor interactions identifies deficient semaphorin signaling between abCaps and the alveolar niche in BPD and BPD+PH

To characterize how abCaps might interact with other cells in the alveolar niche, we interrogated our scRNAseq data with CellChat, which predicts signaling interactions by expression of ligand-receptor pairs. In intracellular communication, cells can be classified as having more outgoing or incoming signaling information^16^. Broadly, CellChat identified endothelial capillaries as acting predominantly as “receivers” of cell-signaling inputs, with AT1 cells and mesenchymal cells having a higher “sender” score (**Fig. 3A**). CellChat identified semaphorin signaling as a leading differentially expressed candidate pathway differentiating term controls and infants with BPD and BPD+PH, with significantly decreased interactions in both epithelial-capillary and mesenchymal-capillary interactions (**Fig. 3B**). Interrogation of the transcriptomic dataset for the specific ligands and receptors of this pathway revealed that BPD+PH patients had markedly decreased expression of *SEMA3B* by AT1 cells, *SEMA3C* by alveolar and ductal myofibroblasts, and *SEMA6A* by endothelial capillary cells, with some decreased expression of these ligands in patients with severe BPD without PH, compared to controls (**Fig. 3C**). Expression of semaphorin receptors *NRP1* and *PLXNA2* was also decreased (**Fig. 3C**) in BPD and BPD+PH patients. Some of these predicted expression changes were spatially validated by RNA ISH, showing decreased expression of *SEMA3B* in *AGER*-expressing AT1 cells and decreased *SEMA6A* in *PECAM1*+ alveolar endothelial cells (**Fig. 3D-E**).

**Fig. 3:**
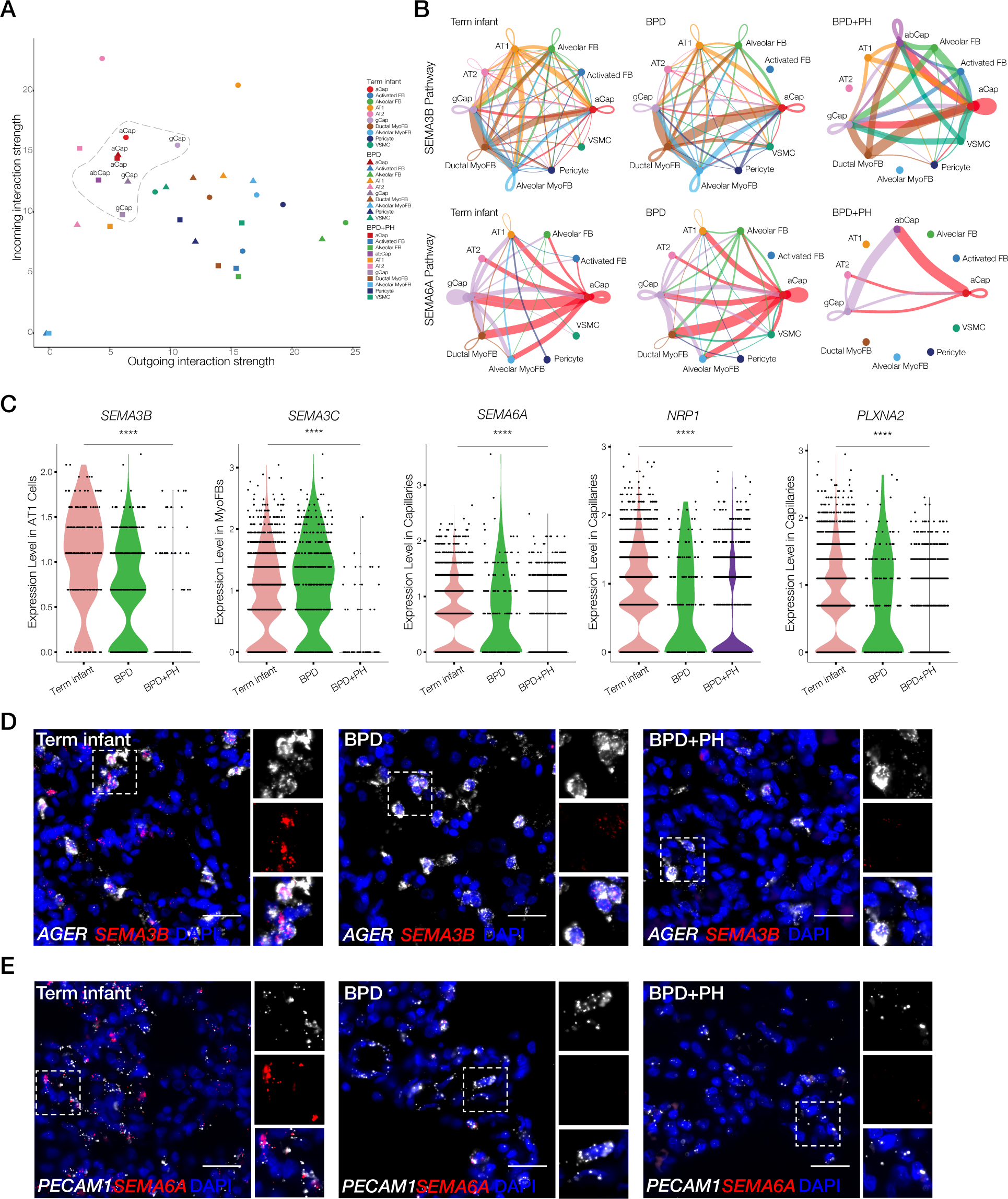
Analysis of predicted ligand-receptor interactions by single-cell sequencing identifies decreased semaphorin signaling between endothelial cells and alveolar niche in the setting of BPD and BPD + PH. A) Cell types plotted by predicted outgoing vs incoming information averaged for all detected pathway interactions from the CellChat ligand and receptor database. B) Circle plots for the SEMA3 and SEMA6 pathways displayed by disease state. The line weights indicate signal interaction strength from gene expression of ligands and receptors for each respective pathway. C) Violin plots for expression levels of semaphorin pathway genes *SEMA3B*, *SEMA3C*, *SEMA6A*, *NRP1*, and *PLXNA2* by disease state, with cell-type specificity indicated on y-axis. **** *p*-value<0.0001 by one-way ANOVA. D) RNA ISH for *SEMA3B* (red) and *AGER* (white), demonstrating co-localization of *SEMA3B* within *AGER*+ AT1 cells in term controls, with decreased expression in patients with BPD and BPD+PH. Scale bar = 25 μm. E) RNA ISH for *SEMA6A* (red) and *PECAM1* (white), demonstrating co-localization of *SEMA6A* within *PECAM1+* endothelial cells in term controls, with decreased expression in patients with BPD and BPD+PH. Scale bar = 25 μm.

### Decreased semaphorin ligand expression replicated in a murine BPD model

To determine whether the decreased semaphorin signaling seen in human tissues could be modeled *in vivo*, we examined a murine model of saccular-stage injury for semaphorin ligand expression. Injury by exposure to hyperoxia (70% O_2_) and inflammation (by intranasal LPS) in the saccular stage (P1-P5) mimics many of the features of BPD, including impaired alveologenesis (as measured by increased mean linear intercept, a measurement of terminal airspace subdivision) (**Fig. 4A-C**)^17^. RNA ISH on samples from this model demonstrated decreased expression of semaphorin ligands *Sema3b* and *Sema6a* in a cell-type specific pattern (**Fig. 4D-E**) similar to that observed in human patients (**Fig. 3D-E**). Notably, by RNA ISH, there was abundant expression of *Sema3b* and *Sema6a* in uninjured AT1 cells and endothelial cells, respectively, suggesting a conserved role of semaphorin signaling in early alveologenesis under normal developmental conditions.

**Fig. 4:**
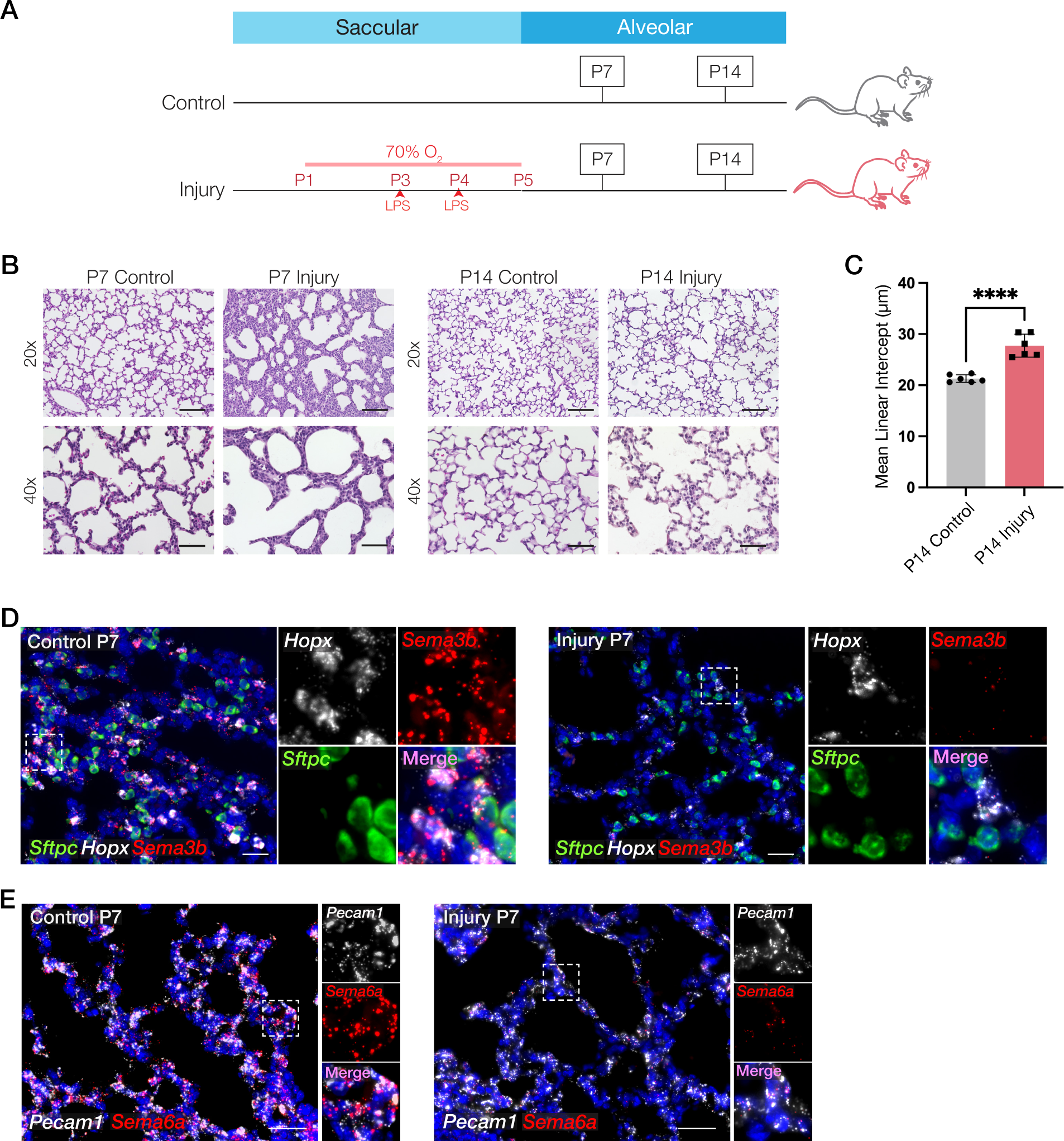
Murine mouse injury models of BPD and BPD+PH display decreased semaphorin signaling. A) Schematic of murine BPD model with hyperoxia exposure from P1-5 and intranasal LPS on P3 and 4. B) H&E of the BPD murine mouse injury model and control mice at P7 and P14 at 20x and 40x magnification. Scale bar = 110 μm (20x), 55 μm (40x). C) Mean linear intercept on the P14 control and injured mouse H&E images at 40x magnification, **** *p-*value<0.0001 by Welch’s *t*-test. D) RNA ISH of *Sftpc* (green), *Hopx* (white), and *Sema3b* (red) on the P7 control and injured mouse lungs exhibiting co-localization of *Sema3b*+ within *Hopx*+ epithelial cells in the alveolar parenchyma. Scale bar = 20 μm. E) RNA ISH of *Pecam1* (white) and *Sema6a* (red) on the P7 control and injured mouse lungs exhibiting co-localization of *Sema6a*+ within *Pecam1*+ endothelial cells in the alveolar parenchyma. Scale bar = 35 μm.

### Human infants with ACDMPV demonstrate decreased semaphorin ligand and receptor expression

Previous work has linked semaphorin signaling to regulation by endothelial-cell transcription factor FOXF1^18^, a known upstream mediator of angiogenesis, and mutations in *FOXF1* result in alveolar capillary dysplasia with misalignment of pulmonary veins (ACDMPV), a rare and lethal form of developmental lung disease with severe pulmonary hypertension. To further explore the relationship between FOXF1 and semaphorin signaling, we interrogated a recently published single-nucleus RNA sequencing dataset from infants and children with ACDMPV and age-matched controls^11^ (**Fig. 5A, S2**) for expression of semaphorin ligands and receptors. While the expression deficits in semaphorin signaling are not identical in ACDMPV and BPD+PH (**Fig. 5B**), we note that patients with ACDMPV have decreased *SEMA3B* expression in AT1 cells and decreased *SEMA6A* signaling in alveolar capillaries (aCaps + gCaps), reminiscent of the deficits in BPD+PH.

**Fig. 5:**
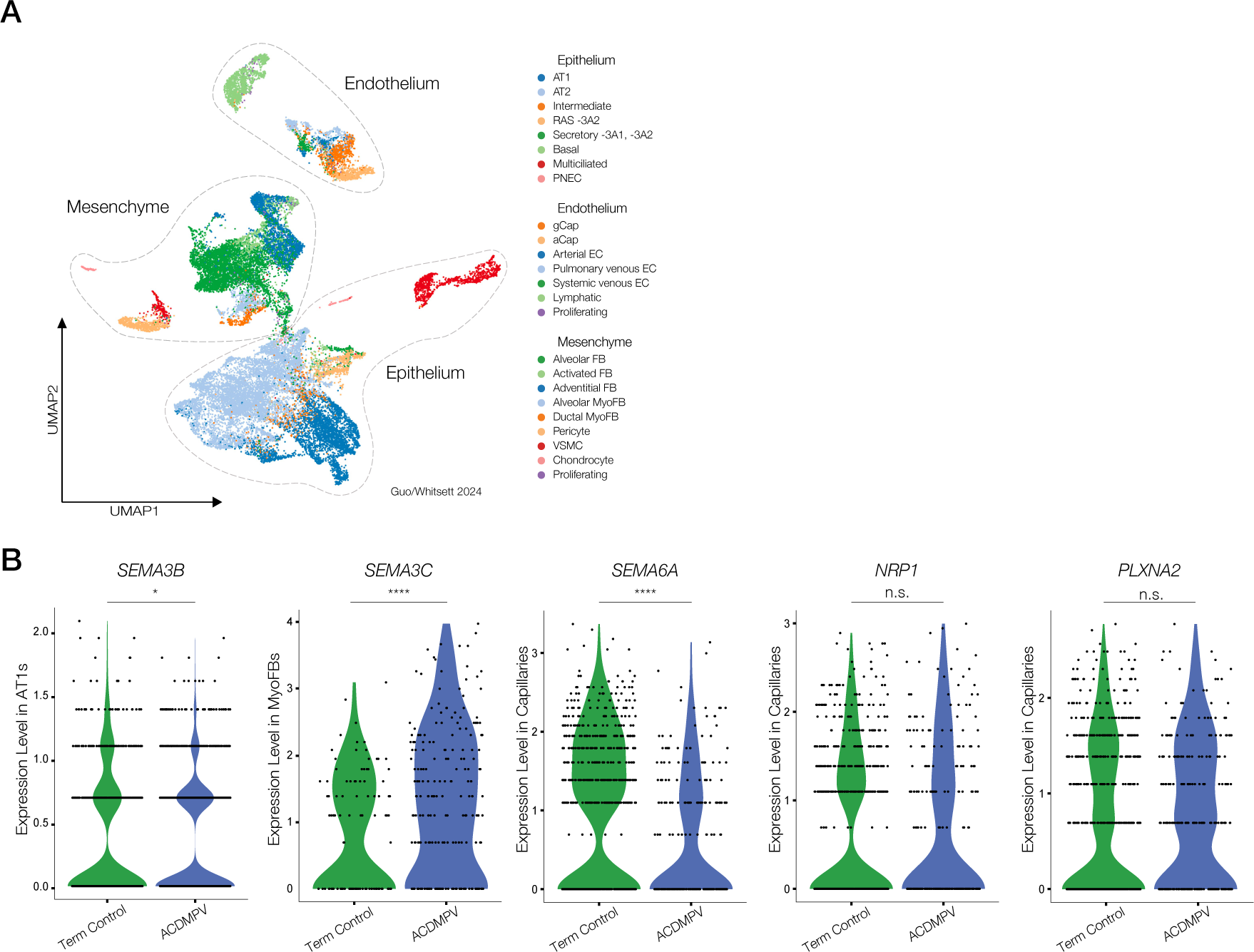
Analysis of human infant ACDMPV dataset identifies decreased semaphorin signaling. A) UMAP embedding of single-nucleus RNA sequencing of five ACDMPV neonatal lung samples^11^, three preterm, and three term control lung samples with an age range of 29 weeks gestational age to 3.5 years old annotated by lineage and cell type. B) Violin plots for expression levels of semaphorin pathway genes *SEMA3B*, *SEMA3C*, *SEMA6A*, *NRP1*, and *PLXNA2* by disease state, with cell-type specificity indicated on y-axis. * *p*-value<0.05, **** *p-* value<0.0001 by Welch’s *t*-test.

### BPD and BPD+PH lung tissues demonstrate decreased FOXF1 expression

Interrogation of our neonatal transcriptomic dataset demonstrated significantly decreased *FOXF1* in the capillary endothelium in BPD and BPD+PH conditions (**Fig. 6A**). Immunofluorescence for FOXF1 and the capillary hallmark protein ERG on BPD, BPD+PH, and term infant control lung tissue substantiates the decreased nuclear localization of FOXF1 within endothelial cells of both disease states (**Fig. 6B-D**), suggesting a functional deficit of this critical transcription factor in BPD and BPD+PH.

**Fig. 6:**
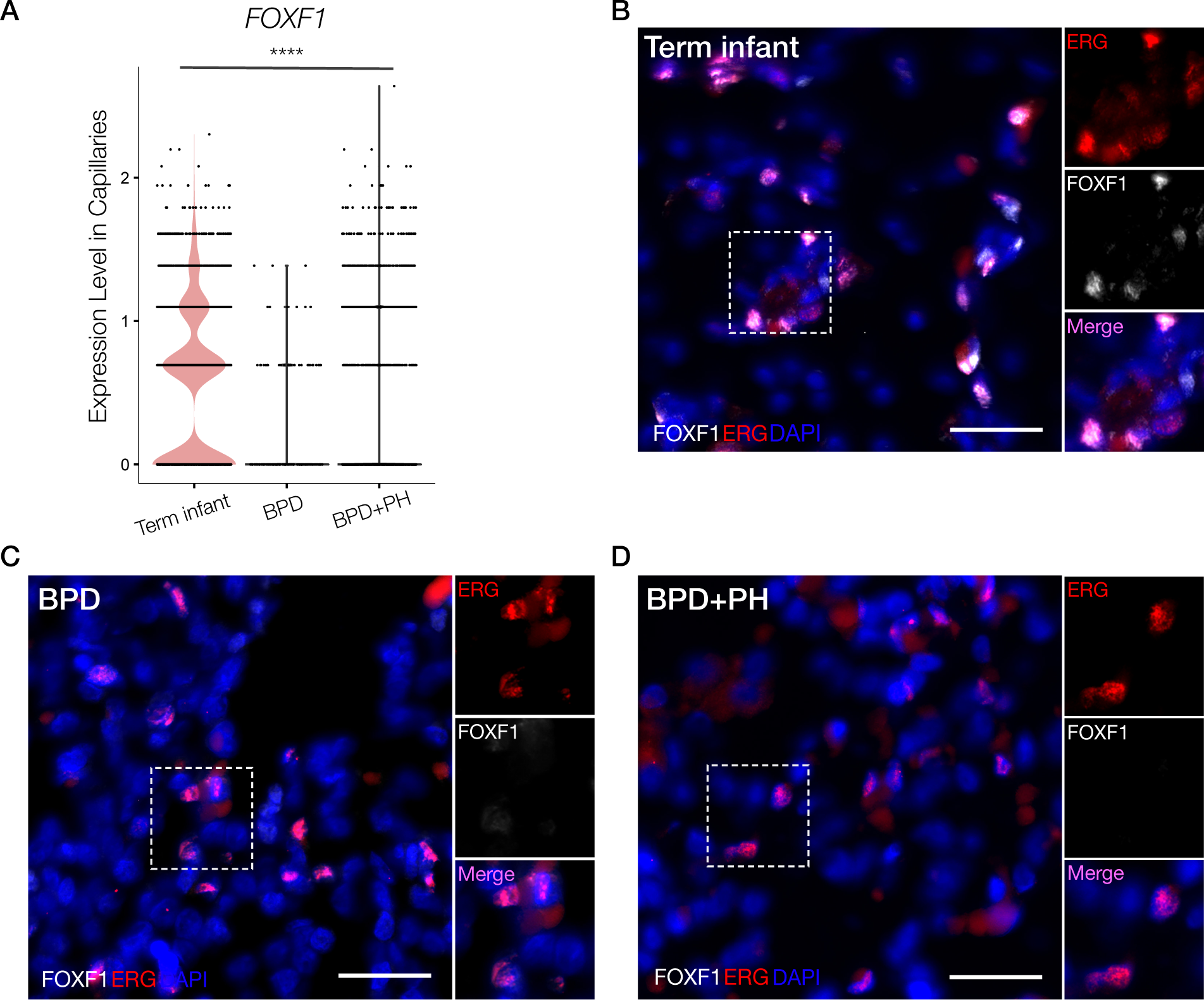
FOXF1 has decreased expression in human neonatal lung injury. A) Violin plots for expression levels of endothelial transcription factor *FOXF1* in the human neonatal BPD and BPD+PH single-cell RNA sequencing dataset comparing disease state in capillaries, **** *p*-value<0.0001 by one-way ANOVA. B) RNA ISH for ERG (red) and FOXF1 (white) demonstrating co-localization of FOXF1 within ERG+ endothelial cells in the distal lung parenchyma in term controls, with decreased expression of FOXF1 in patients with (C) BPD and (D) BPD+PH. Scale bar = 25 μm.

## Discussion

In this single-cell transcriptomic analysis of human neonatal lungs across stages of injury, we identified a previously undefined endothelial cell state, abCap, that is enriched in lungs from infants with bronchopulmonary dysplasia and pulmonary hypertension (BPD+PH). Using predictive analysis tools and tissue validation, we discovered deficits in semaphorin signaling and an apparent functional decrease in FOXF1 in infant tissue representing BPD and BPD+PH. The latter suggests a possible link between the rare disease ACDMPV and the severe forms of abnormal lung development that occur with preterm birth. The concordance of these findings in a murine model of BPD provide foundational data that support a new role for semaphorin signaling as a critical mediator of normal lung development whose disruption may lead to the abnormal alveologenesis and vascular development observed after preterm birth^19^.

There are potential limitations in this study. While transcriptomic data from these neonates at varying stages of development and evolving injury are informative in generating new hypotheses about molecular mechanisms that drive abnormal lung development after preterm birth, they come from a limited number of subjects, and autopsy samples in general have an inherent bias toward the sickest individuals who have died from BPD and BPD+PH. Nonetheless, these data fill a critical gap regarding the deep transcriptomic analysis of normal and injured lungs across the lifespan, because there are so very few published reports on neonatal premature injured and uninjured lung samples.

A central strength of our study is the transcriptomic integration of these rare samples with samples from ACDMPV patients. The observed deficits in semaphorin signaling common to BPD+PH and ACDMPV suggest that the functional deficits in *FOXF1* associated with the developmental phenotypes of BPD and BPD+PH may be mechanistically related to the observed deficits in ACDMPV. Emerging technologies in spatial transcriptomic analysis offer increasing potential for interrogation of archival FFPE tissue blocks, which we anticipate will facilitate rapid growth in studies across the neonatal and pediatric populations. Making these emergent data accessible and searchable could promote discovery of plausible mechanistic targets for promoting lung growth and regeneration and allow for a larger representation of samples across institutions and platforms.

While further work is required to define when abCaps emerge in disease and the role they may play in the pathogenesis of BPD, the discovery and validation of the abCap cell-state broadens our understanding of the endothelium as a receiver and coordinator of cell signaling and cell movements during this specific developmental stage. Analysis across hundreds of published datasets reveals that abCaps are distinct from the diversity of endothelial cell types and states observed in other diseases (e.g. adult PH, COVID, ARDS), and suggests that abCaps may be a cell state specific to BPD+PH. Deeper characterization of these cells demonstrates increased expression of YAP signaling targets^20–22^ (e.g., *CCN1, CCN2, SERPINE*1, and *CRIM1*, along with abCap hallmark and YAP target gene *ANKRD1*). While the VEGF-induced YAP/TAZ signaling responsiveness in endothelial cells is required for vasculogenesis^20-23^, elevated YAP/TAZ target expression in abCaps in BPD+PH patients could indicate an attempted cell-autonomous compensatory response within abCaps themselves or arising as a response to abnormal cell-signaling by injury-affected adjacent cells. Indeed, in murine models of hyperoxia-induced injury in neonates and hypoxia-induced PH in adults, an increase in endothelial CCN gene expression and *Ankrd1* expression was reported^24–26^. Understanding how failed injury repair after preterm birth leads to severe BPD+PH, and how abCaps affect and respond to the complex signaling milieu of the post-injury alveolar niche is critical to untangling the mechanisms of abnormal repair in BPD and BPD+PH^27^.

CellChat analysis predicted alveolar capillaries as receivers of signaling information from adjacent alveolar epithelial and mesenchymal cells. In BPD and BPD+PH, and murine hyperoxia BPD models, we noted dramatic reductions in expression of factors involved in semaphorin signaling. The semaphorin pathway is well defined in neuronal axon movement and synapse coordination, especially in the cellular targeting of growth cones toward target cells^28^. While the role of this pathway in lung development has generally received little attention, a global Sema3a inactivation resulted in high perinatal lethality with marked impairment in alveologenesis in rare surviving KO pups^29^, global knockdown of *Sema3C* impaired alveologenesis^30^, and ectopic *Sema3a* overexpression attenuates apoptosis in neonatal rats exposed to hyperoxia^31^. Further, genetic inactivation of semaphorin receptor Nrp1 (neuropilin-1) in the adult mouse impairs airspace remodeling after smoke exposure^32^, suggesting a role for semaphorins in repair after injury. This study builds from these foundational papers to dissect the cell-type specific expression of distinct semaphorins in the setting of alveologenesis and injury-repair. Our findings on semaphorins add to previous work identifying axonal-guidance signaling among several pathways differentially methylated in BPD^33^ as well as those made on another axon-guidance pathway (Netrin/DCC/Unc) in fine-tuning the size and shape of emerging epithelial buds during early lung development^34^, although the latter study is of a far earlier developmental stages than the alveolar stage reported here.

A central role of semaphorins is in regulating movement of cells toward or away from each other. Recent 4D-live imaging experiments suggest that alveologenesis results from precisely coordinated migration and shape change of epithelial and endothelial cells as they move away from the alveolar duct to form a common functional unit of gas exchange^35^ with AT1 and aCap cells tightly aligned on a shared basement membrane, and our data strongly implicate the semaphorin pathway as a regulator of these complex cellular movements and shape changes required for alveolar formation. In BPD+PH, whether abCaps still positively contribute to alveolar function or are the major contributor to deficit is unclear. Although abCaps present with characteristics notably distinct from prior published cell populations, they do share some transcriptional homology with endothelial tip cells, a specialized epithelial cell that provides guidance for the growth and migration of capillaries during angiogenesis.

Expressing some but not all features of endothelial tip cells in evolving BPD+PH may be indicative of compensatory response during repair or representative of a misaligned developmental program in abCaps as a result of decreased cues from semaphorins and other intercellular signals. Together, the decreased expression of *SEMA3B* by AT1 cells, *SEMA6A* by aCaps and gCaps, *SEMA3C* in the mesenchyme and decreased *NRP1* expression in endothelial cells under arrested lung development brings semaphorin signaling to the foreground in the injured and normal contexts. A deeper characterization of the timing and cell specificity of semaphorin signaling during lung development may reveal critical windows in which this pathway could be targeted to promote regrowth and structural repair after injury.

## Methods

### Human Infant Lung Repository

At the time of autopsy/biopsy, fresh lung tissue was isolated and processed for single-cell sequencing (scRNAseq) as described previously^5^ with additional details below. In addition, we studied formalin-fixed paraffin embedded tissue blocks from human infant lung samples from autopsies and biopsies in varying stages of lung injury and without evidence of lung injury (as determined by a clinical pathologist) have been collected at Vanderbilt University Medical Center. All samples are de-identified with the exception of sex, gestational age at birth, age at time of death, cause of death, and whether or not the patient had evidence of BPD (defined as a need for supplemental oxygen at 36 weeks correct gestational age) or pulmonary hypertension (as determined by clinical echocardiography). A table with the clinical characteristics of the infant lungs used for FFPE tissue and scRNAseq is found in the supplement (**Supplementary Tables 1-2**). This infant lung repository has been reviewed and approved by the Vanderbilt University Institutional Review Board.

### Single-cell sequencing

Human lung tissue from infants who died with BPD, BPD+PH, acute preterm lung injury, and term infant controls were processed into single-cell suspension as described previously^5^. scRNAseq libraries were generated using the 10x Chromium platform 3’ or 5’ library preparation kits (10x Genomics) following the manufacturer’s recommendations and targeting 10,000-20,000 cells per sample. Next-generation sequencing was performed using an Illumina Novaseq 6000. Reads with read quality less than 30 were filtered out and CellRanger Count v7.1.0 (10x Genomics) was used to align reads onto the GRCh38 reference genome. Full code used in the analysis of these data is available at https://github.com/SucreLab/HumanNeonatal. A table with the clinical characteristics of infants whose lungs were sequenced is in the supplement (**Supplementary Table 1**).

### Single-cell RNA sequencing analysis

The ambient RNA was removed from the CellRanger count matrix h5 file outputs using CellBender v0.2.1^36^. The scRNAseq data was analyzed using Seurat (v4.4.0)^37^ in R (v4.3.2). The quality control of the ambient RNA corrected cells filtered out cells with >10 % mitochondrial mRNA mapped UMIs, <500 identified genes, and >7000 identified genes. The data was normalized using SCTransform (v0.4.1)^38^ with the glmGamPoi (v1.14.0)^39^ method by stabilizing the variance of the total molecular counts. The default parameters were used with SCTranform except for: batchvar set to correct batch effects from the tissue samples processed, vars.to.regress set to regress out the effects on differential gene expression analysis due to the presence of increased mitochondrial mRNAs and ribosomal rRNAs, and ncells set to NULL. The reduction of dimensionality was calculated via PCA. The PCA embeddings were further integrated with Harmony (v1.2.0)^40^ to account for any remaining batch effects from the samples. The Harmony reduction was then applied in obtaining the UMAP embeddings and the shared nearest neighbor graph which subsequently allowed for the Louvain algorithm to find the cell clusters.

The clusters were split into the four main cell subtypes of epithelial, endothelial, mesenchymal, and immune based on respective hallmark genes (**Supplementary Table 3**). Each subtype went through an iterative process of SCTransform, PCA, Harmony, UMAP, shared nearest neighbors, and clustering to isolate and remove any remaining doublet or nonbiological cells. For each iteration, the cluster resolution would start in the range of 1 < n <= 2, and with each iteration, n would be brought down slightly until the last iteration which would have n = 1. Cell type labels were assigned after quality control and cleaning of the data from their respective hallmark genes (**Supplementary Tables 3-4**). After annotation, the four subtype-group Seurat objects were merged to produce the final Seurat object.

CellChat^16^ was used to predict possible ligand-receptor pair interactions between cell types of the same condition to generate hypotheses about possible signaling mechanisms associated with BPD and BPD+PH. We employed SCimilarity^15^ to query cell populations or states that could have similar gene expression as the BPD+PH endothelial cell state (abCaps). To do this, we used a gene signature list curated from the intersection of the top Wilcoxon rank sum method ranked genes, top log2 fold change expression genes, and genes with the highest proportion expressed within abCaps versus all other cell types: *ANKRD1, CCN2, KITLG, SERPINE1, TNFRSF12A, CRIM1, GDF15, CCND1, EDN1, RBFOX2, CD151, PLS3, CD59, GBE1, PODXL*. We then applied this gene module score to all cell populations or clusters in each study independently, with a cutoff of a cell score greater than two standard deviations from the mean cell score in the dataset. In summary, we looked for cells with features of the expression pattern identified in the abCap gene module. We then plotted the percentage of cells within each cluster that passed this cutoff.

Reanalysis of the recently published single-nucleus RNA sequencing dataset of infants who died of ACDMPV and term control infants was performed using the same strategy used above for scRNAseq.

### Murine model of neonatal lung injury

C57BL/6 mice were exposed to 70% O_2_ (hyperoxia) or 21% O_2_ normoxia conditions from P1 until 5 days of age (P5), with hyperoxia animals receiving intranasal LPS at 1 µg/g on P3 and P4. On P7 and P14, lungs were inflation-fixed by gravity filling with 10% buffered formalin and paraffin embedded as described previously^41^. This protocol was approved by the Institutional Animal Care and Use Committee of Vanderbilt University and was in compliance with the Public Health Services policy on humane care and use of laboratory animals.

### RNA in situ hybridization

RNAScope technology (ACDBio) was used to perform all RNA ISH experiments according to the manufacturer’s instructions as described previously. RNAScope probes to the following human genes were used (*SEMA3B, AGER, SFTPC, SEMA6A, PECAM1).* RNAScope probes to the following mouse genes were used*: Sftpc, Hopx, Pecam1, Sema3b, Sema6a.* Positive control probe (PPIB) and negative control probe (DapB) were purchased from ACDBio and performed with each experimental replicate. When RNAScope and IF are performed together, this was done in accordance with the manufacturer’s instructions.

### Immunofluorescence

Immunofluorescence was performed as described previously, with the following primary antibodies used: anti-PDPN (DSHB, Iowa), anti-FOXF1 (1:100, R&D systems AF4798), anti-ERG1 (1:100, Abcam ab214341) and the following secondary antibodies used: 1:200 Donkey anti-Goat (Invitrogen A21447), 1:200 Donkey anti-Mouse (Invitrogen A31570).

### Image acquisition and analysis

We acquired fluorescent and brightfield images using a Keyence BZ-X810 with BZ-X Viewer software and 40× objective. Images were collected with the following filter sets: 405, 488, 561 and 647 nm. Automated image analysis was performed with HALO software (Indica Labs) as described previously. After HALO analysis, the rate of false co-localization of non-physiological expression combinations (*Sftpc+Pecam1+*) was low, with a mean of <1%.

### Lung Morphometry

Mean linear intercept was performed on a minimum of 10 40x images per mouse (n=6 mice per group) as described previously ^42^, with the use of ImageJ and AlveolEye for automated calculation of MLI measurements (https://github.com/SucreLab/AlveolEye).

### Statistics

Statistical analysis was performed in R and with Prism Graphpad version 10.0.03, with specific details on statistical tests provided in Fig. legends.

## Data Availability

The single-cell sequencing data generated for this manuscript are available as a freely searchable tool at www.sucrelab.org/lungcells. As noted above, all code used in analysis and generation of sequencing data display can be found at https://github.com/SucreLab/HumanNeonatal. Raw sequencing data will be deposited at GEO and available for download. We will eagerly provide processed data and objects upon request.

## Acknowledgements

This work was supported by the Francis Family Foundation (JAK, DBF, JMSS, NMN), R01HL168556 (JMSS), K08HL143051 (JMSS) R01HL145372(JAK/NEB), R56HL167937 (DBF), U01HL175444 (JAK, NEB, JMSS). We are grateful to Anne Hilgendorff and Vivian Siegel for thoughtful discussions and insights.

## Author Contribution

Conceptualization: JAK, JMSS, DBF, CVEW, Methodology: SPS, JAK, NMN, CSJ, ALS, Validation: SG, GF, DW, Resources: MEK, DBF, Investigation and Interpretation: CSJ, JAK, JMSS, SPS, NMN, BDF, CVEW, SM, NEB Manuscript preparation (original): SPS, JMSS, Manscript preparation (revision): SPS, NMN, CSJ, ALS, SG, MEK, GF, SM, NEB< CVEW, DBF, JAK, JMSS.

---

**Supplementary Table 1:** Clinical characteristics of infants whose lung tissue samples were analyzed by single-cell RNA sequencing, CGA = corrected gestational age.

| Gestational Age | Age at time of sample collection | Autopsy/Biopsy | Clinical Metadata |
| --- | --- | --- | --- |
| 40 wks | 7 days | Biopsy | Normal tissue from infant s/p CPAM resection |
| 37 wks | 20 days | Biopsy | Normal tissue from infant s/p CPAM resection |
| 27 wks | 7 months<br>(52 wks CGA) | Autopsy | BPD + PH |
| 26 wks | 7 months<br>(51 wks CGA) | Autopsy | BPD + PH |
| 27 wks | 7 months<br>(52 wks CGA) | Autopsy | BPD |
| 25 wks | 7 months<br>(50 wks CGA) | Autopsy | BPD |
| 26 wks | 2 wks<br>(28 wks CGA) | Autopsy | Intubated 14 days; Died with severe IVH |

**Supplementary Table 2:** Clinical characteristics of infants with lung tissue samples in blocks that were studied by RNA *in situ* hybridization and immunofluorescence.

| Gestational Age | Age at time of sample collection | Autopsy/Biopsy | Clinical Metadata |
| --- | --- | --- | --- |
| 35 wks | 2 days | Autopsy | HIE |
| 39 wks | 3 days | Autopsy | HIE |
| 39 wks | 1 day | Autopsy | Stillbirth |
| 24 wks | 4 months | Autopsy | BPD |
| 26 wks | 6 months | Autopsy | BPD |
| 27 wks | 7 months | Autopsy | BPD + PH |
| 26 wks | 7 months | Autopsy | BPD + PH |
| 27 wks | 4 months | Autopsy | BPD + PH |

**Supplemental Figure 1:**
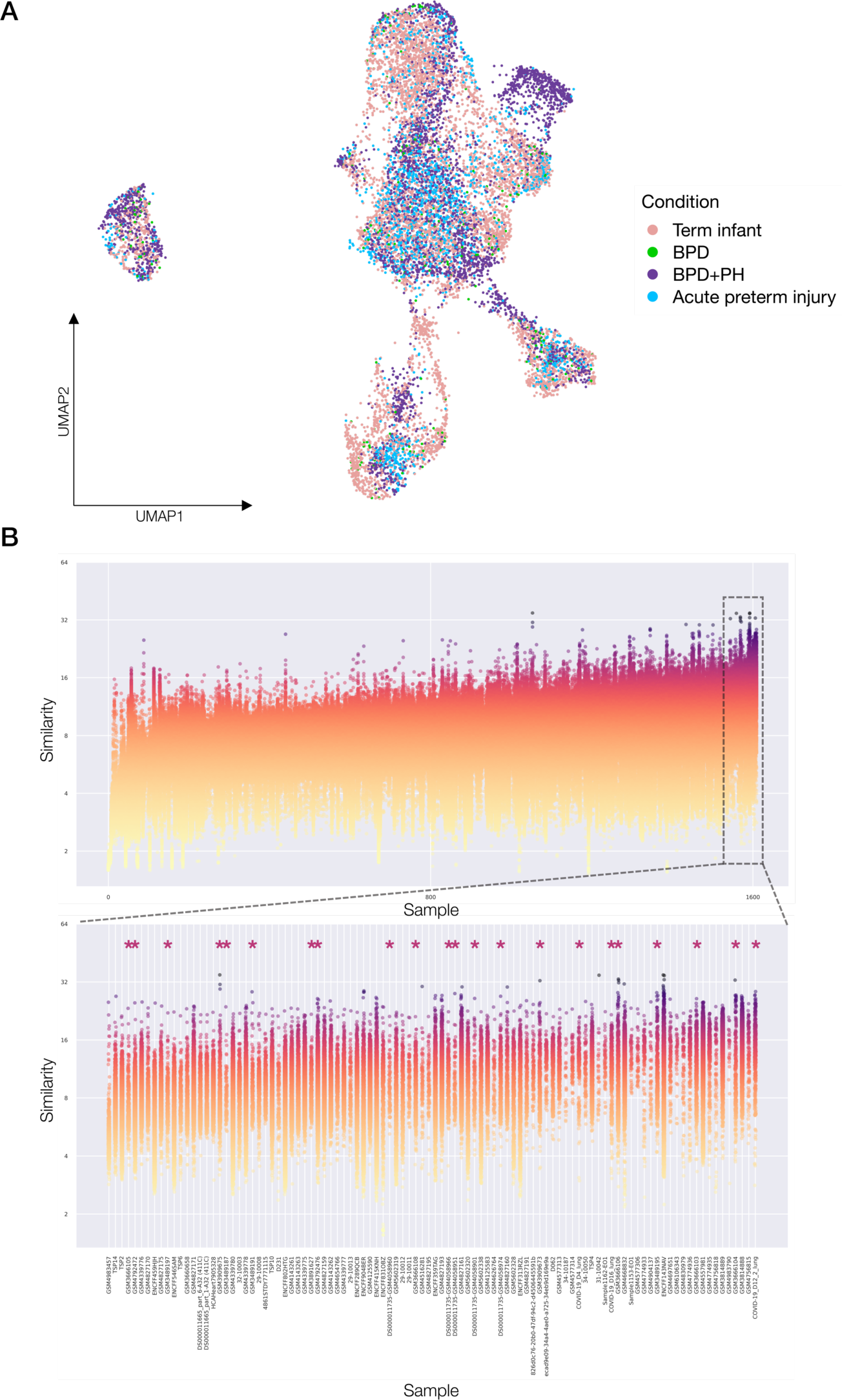
A) UMAP embedding of the neonatal human lung endothelial cells labeled by disease-state condition. B) 1,610 identified samples containing endothelial cells filtered by biological factors and by top 100 SCimilarity query score compared to the expression profile of abCaps, identified 22 lung sequencing samples contained within 6 studies of the most comparable cell populations.

**Supplemental Figure 2:**
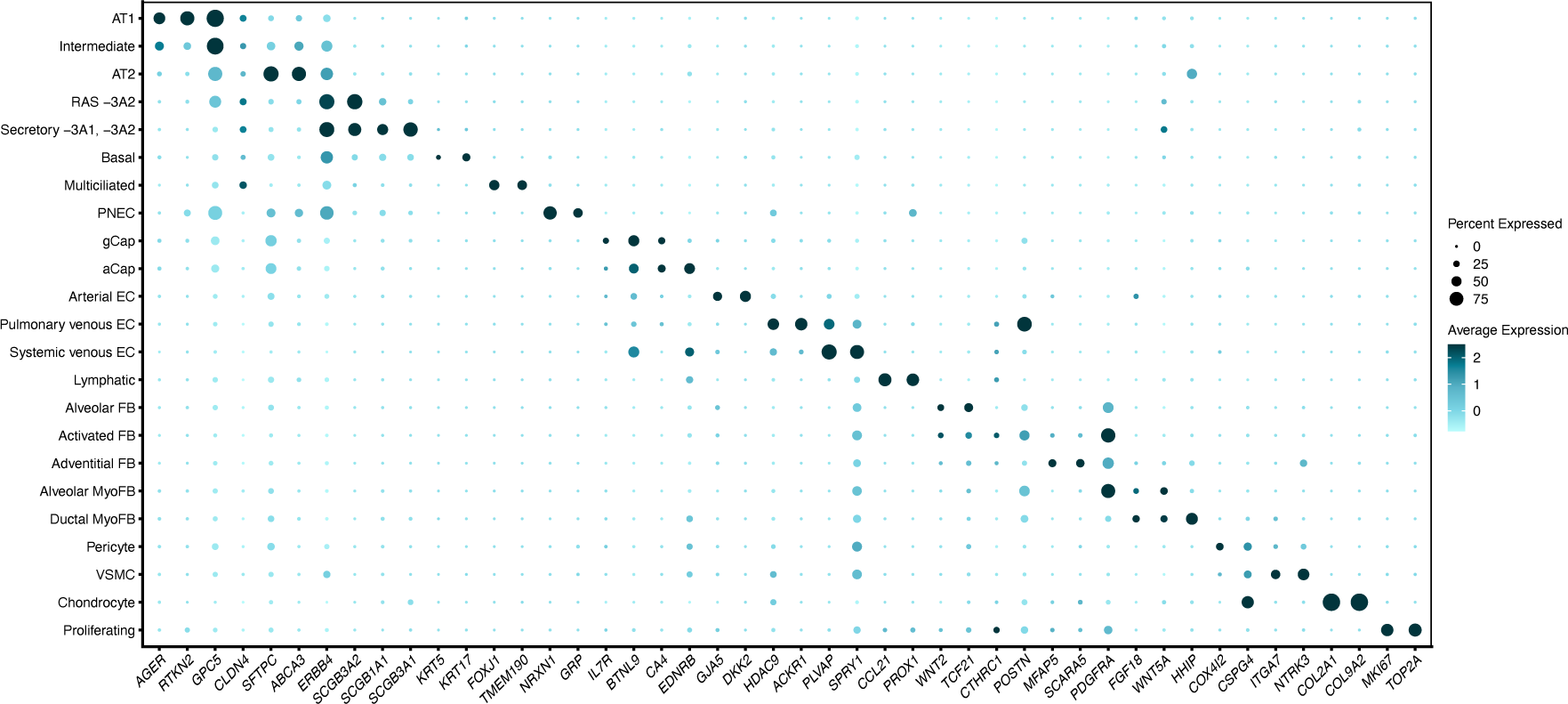
Dot plot of the hallmark-identifying genes for each cell type cluster of the human infant ACDMPV snRNAseq reanalysis dataset. Dot size corresponds to the percentage of cells within a cluster expressing a certain gene. Color intensity corresponds to the relative average expression level.

**Supplementary Table 3:**
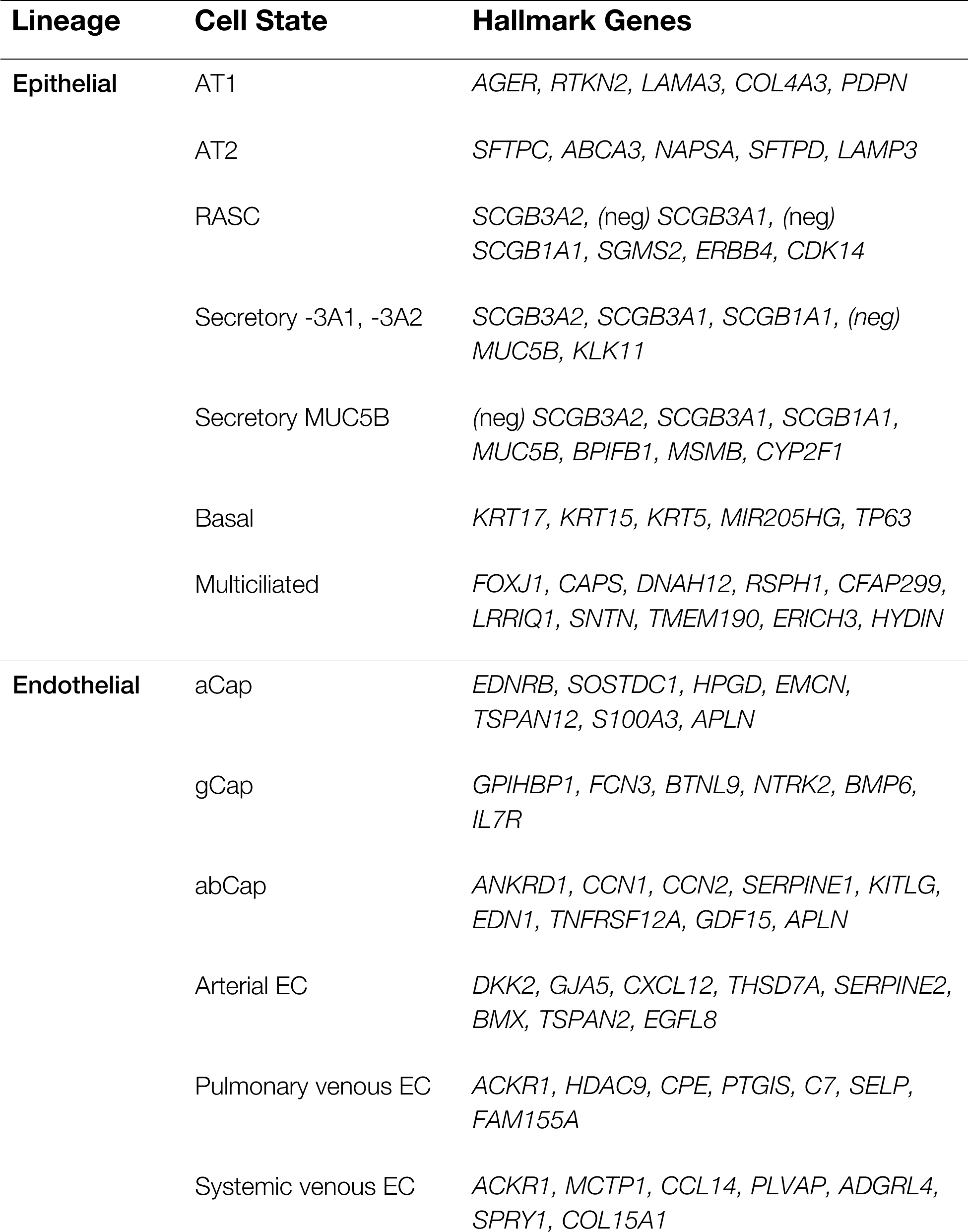
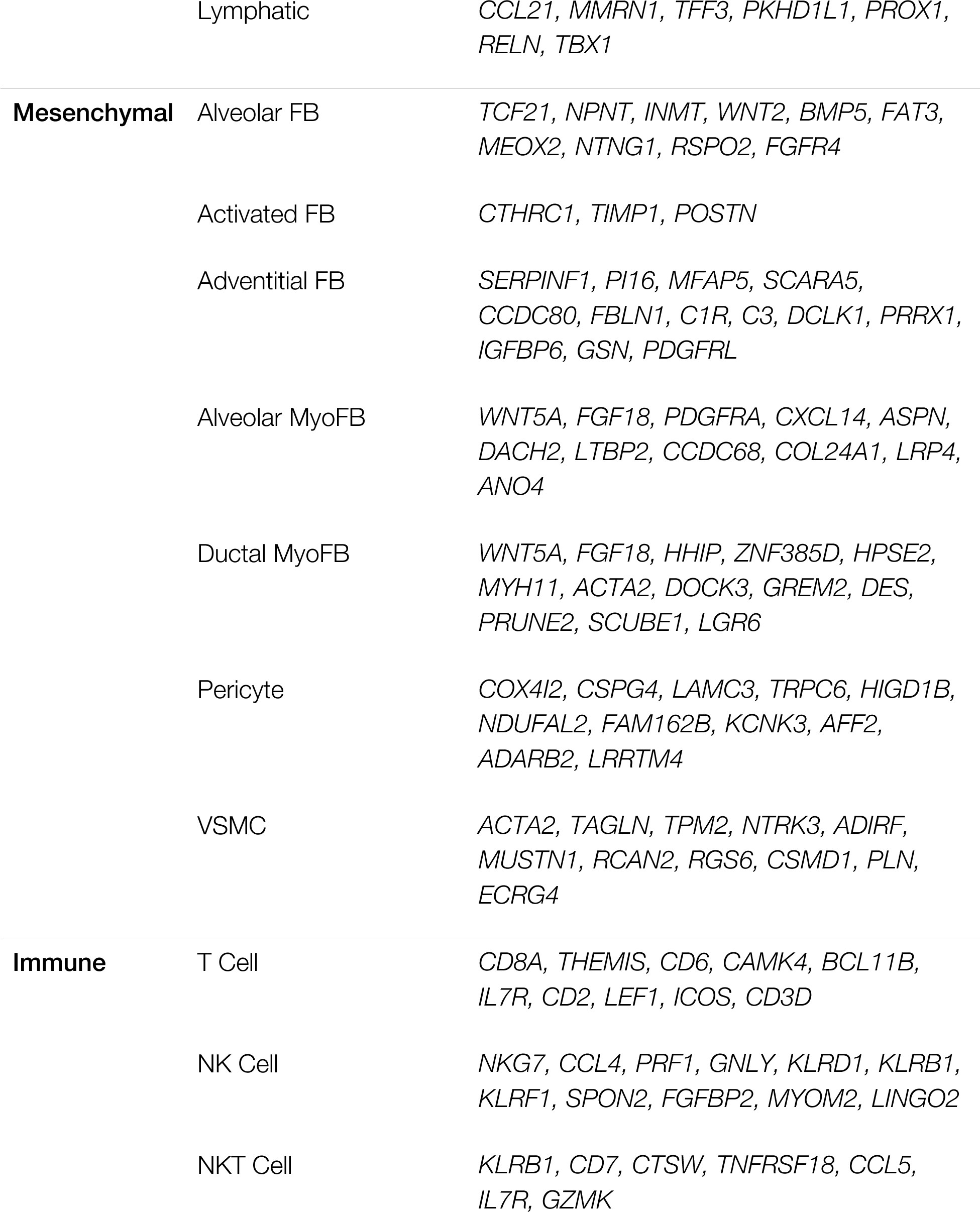
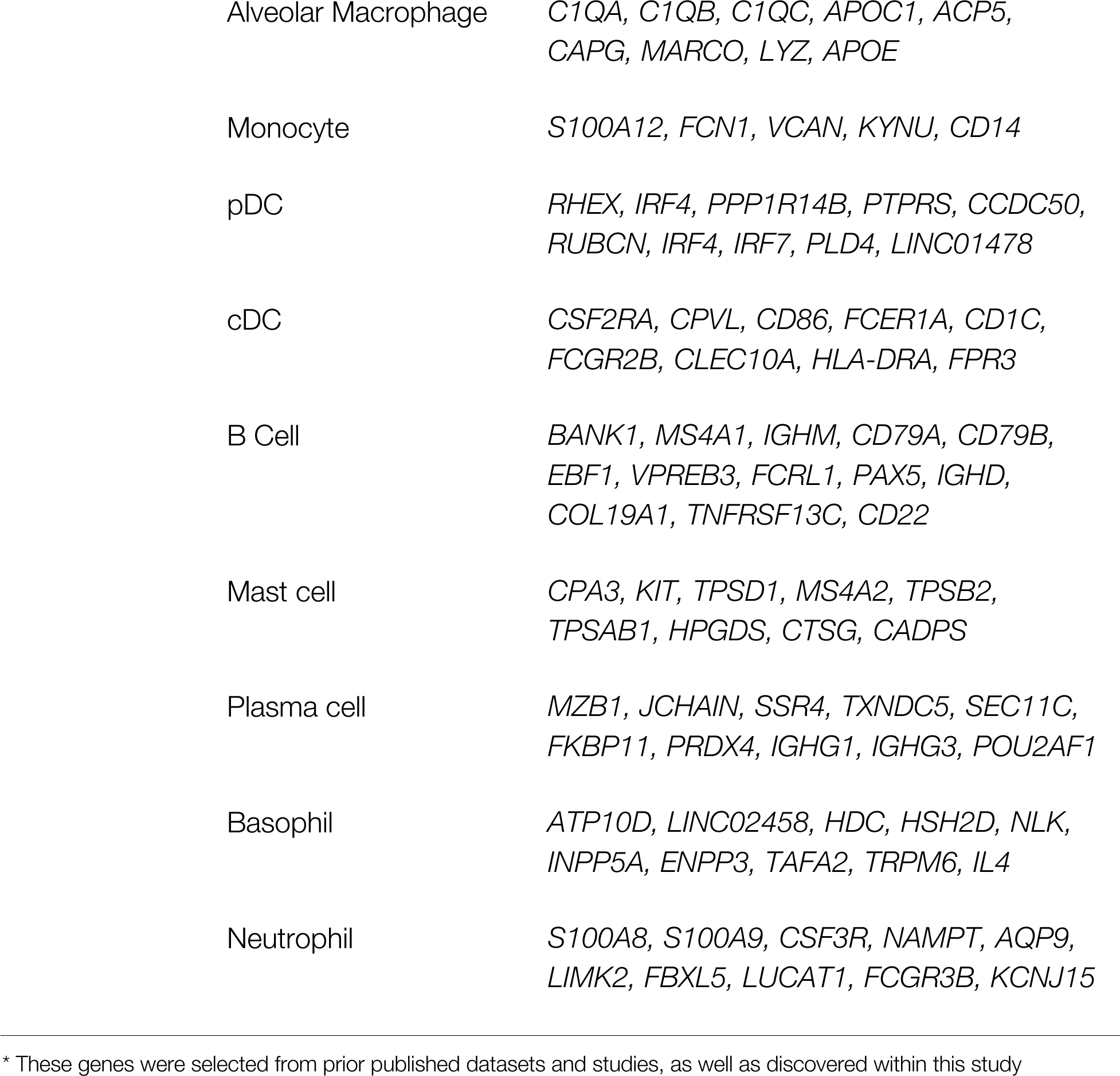
Select hallmark genes of the 33 identified cell type populations.

